# OPM quantum sensors enhance non-invasive neuroimaging performance

**DOI:** 10.1101/2025.05.30.657044

**Authors:** M. Brickwedde, P. Anders, P. Krüger, T. Sander, P. J. Uhlhaas

## Abstract

Much of our understanding of how neural circuit activity relates to human behaviour as well as mental health stems from non-invasive assessments of the central nervous system. Further progress in key questions, however, requires a higher differentiation of circuit-level activity, which remains constrained by the sensitivity and resolution of existing instruments. Here we show that neural quantum sensing with optically pumped magnetometers (OPMs) addresses this issue by significantly outperforming EEG and conventional MEG in both signal-to-noise ratio (SNR) and signal coherence across trials. SNR for OPMs was increased by up to 205% compared to EEG and by up to 40% compared to conventional MEG. Likewise, the signal coherence across trials of OPMs was increased by up to 61% compared to EEG and by up to 23% compared to conventional MEG. Our data emphasize the important role of OPMs for non-invasive neuroimaging, paving the way for significant advances in basic research as well as translational applications.

## Introduction

The measurement of neural signals with current non-invasive instruments is constrained by limited temporal and spatial resolution^1^. Accordingly, progress on key questions concerning the human central nervous system is at an impasse, as non-invasive methods lack sufficient performance to advance the differentiation of neural circuit activity, while model organisms often diverge too far from human physiology^1^.

Nearly a decade ago, optically pumped magnetometers (OPMs) have been proposed as quantum sensors for magnetoencephalography (MEG), with the promise of higher signal amplitudes and signal-to-noise ratios (SNRs) compared to the current gold-standard^2–4^. Improvements in these domains could have far-reaching consequences, from a higher differentiability of neural circuit activity^5^, to enhanced precision in individual diagnostics and therapy^6^. Yet despite their potential, a systematic statistical comparison between OPM, EEG and conventional MEG, remains missing. Here we show that SNR and trial coherence of OPMs significantly exceeds EEG and conventional MEG.

Conventional MEG systems rely on superconducting quantum interference devices (SQUIDs), which require continuous helium cooling. As a result, SQUID-sensors can only be accommodated in rigid dewars which place sensors several centimetres away from the human scalp. In contrast, OPMs measure neural signals via light absorption in vaporized alkali atoms, which is modulated by surrounding magnetic fields, such as those generated by neuronal activity^7–9^. Current OPM sensors are compact (volume: ∼5cm^3^)^10^ and can therefore be positioned directly on the human scalp. Because magnetic field strength decays approximately with the square of the distance to the neural sources^11^, OPMs can record neuronal signals with higher signal amplitude ^3^ and detect magnetic fields which may be inaccessible to SQUID sensors. EEG signals, on the other hand, propagate through tissue with varying electrical conductivity - such as the scalp, skull, and cerebrospinal fluid - which leads to spatial smearing and mixing of neural sources at the sensor level^12,13^, thereby reducing measurement precision compared to OPM and SQUID sensors.

Simulations have demonstrated increased signal amplitude, SNR, as well as improved dipole localization accuracy for OPMs compared to SQUID sensors^4,14–17^. Several proof-of-concept studies showed that OPMs measure neural activity such as resting-state connectivity, event-related fields and steady-state responses, which are comparable in pattern and source reconstruction to those measured with SQUIDs^15,18–27^. In direct comparisons, there have been indications that the performance of OPMs superior to EEG- and similar to SQUID-systems, however, contrary results were also reported^15,28–30^. Additionally, comparisons were based on small sample sizes (n < 10)^3,24,31–35^, which limits statistical power and precludes statistical evaluation between measurement modalities. As a result, definitive evidence for the performance of OPMs relative to EEG and SQUID remains missing.

To address this question, we recorded parallel EEG/OPM-data and obtained SQUID-measurements in a separate session within a sample of n = 23 healthy controls and analysed neuronal responses to rhythmic auditory 40 Hz stimulation, an established paradigm in basic as well as clinical research^18,36–39^. Our findings demonstrate that the SNRs and coherence across trials of OPMs significantly exceed EEG and SQUID. It can therefore be expected that OPMs will substantially advance electrophysiological neuroimaging across fields from basic research to precision diagnostic and therapy.

## Results

### Measuring auditory steady-state potentials with OPM, EEG and SQUID

We designed a wearable cap comprising OPM sensor holders and EEG electrodes to allow simultaneous measurement of EEG and OPM (Fig. 1A). To cover right and left temporal areas, we positioned 5 OPM sensors between EEG electrodes on each lateral side of the cap. By choosing 10 SQUID-gradiometers in matching positions to the OPM sensors and 10 EEG electrodes, which best reflected the maximum activity of auditory steady-state responses, we ensured comparability between measurement modalities (see Fig. 1C).

**Figure 1.**
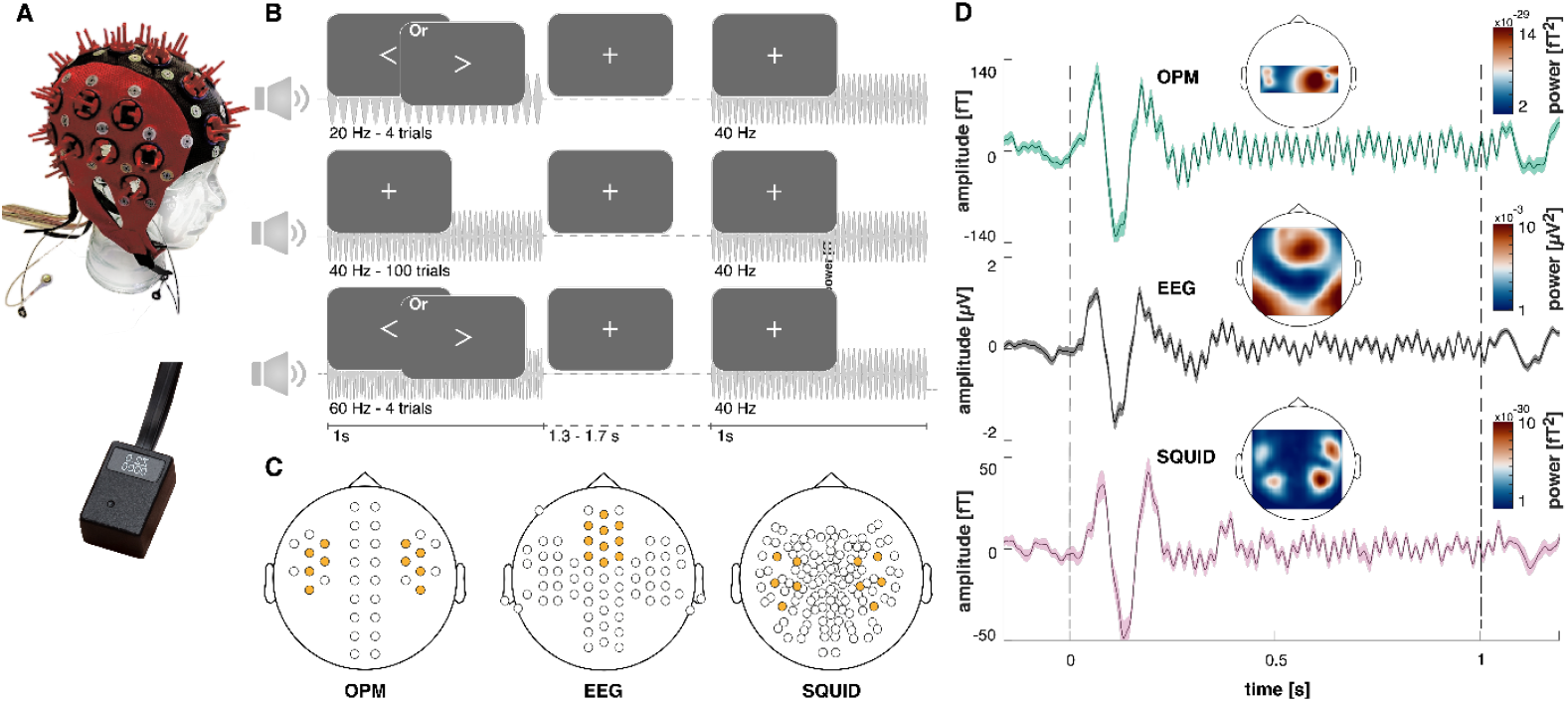
Measuring auditory steady-state potentials with EEG, OPM and SQUID. **A)** EEG and OPM were measured in parallel with a custom cap, which consisted of OPMs (QuSpin, 2^nd^ Gen; https://quspin.com/) inserted into self-made holders in between EEG electrodes. **B)** We played amplitude modulated 40 Hz tones with a duration of 1s and a jittered inter-trial interval of 1.3 to 1.7 s. Participants were asked to fixate to the center of the screen and react to rare stimuli consisting of arrows pointing to the right, while ignoring arrows to the left. In trials with arrows, instead of a 40 Hz tone, either a 60 Hz or a 20 Hz tone were played. **C)** We compared the performance of 10 OPM sensors with 10 EEG electrodes and 10 SQUID gradiometers. Their position (marked in yellow) was chosen to reflect topographic equivalence of the position of SQUID and OPM sensors and, in the case of EEG, the strongest activation of 40 Hz activity. **D)** OPM, EEG and SQUID showed strong correspondence of the measured auditory steady-state response averaged over all participants (n = 23). Displayed is the sensor/electrode with the highest signal-to-noise-ratio for each measurement modality. Due to a reduced distance to the scalp, the signal amplitude was increased by a factor of 2-3 for OPMs compared to SQUID. The topography of the auditory steady-state response in SQUID reflects the dipolar temporal structure expected for auditory responses. The topography for OPMs suggests a similar pattern, however the topography was constrained by the limited number of sensors. The auditory steady-state response of the EEG displayed most strongly over frontocentral electrodes.

To assess measurement precision, we applied 40 Hz rhythmic auditory stimulation, of which the spectral and topographic responses are well-established (Fig. 1B)^18,36–39^. Participants received instructions to focus on the centre of the screen while 40 Hz amplitude-modulated 1s-tones were presented. In order to keep participants engaged, we included a rare go-nogo task, which consisted of either 20 Hz or 60 Hz tones and an arrow, which participants needed to respond to (pointing to the right) or ignore (pointing to the left).

Across measurement modalities, we found a strong similarity in the temporal course of auditory evoked potentials and steady-state responses. As expected, the signal amplitude was increased by a factor of 2-3 for OPM compared to SQUID-signals (see Fig. 1D). The topographic distribution of auditory steady-state responses for EEG was strongest over fronto-central electrodes and over temporal areas for the SQUID measurement, which reflected previous findings^37,38,40^.

### The signal-to-noise ratio of OPMs exceeds EEG and SQUID

We investigated the signal-to-noise ratio (SNR) as peak power in the trial average at 40 Hz in relation to power at adjacent frequencies during the auditory steady-state phase of the stimulation. Additionally, to optimize the steady-state signal representation across sensors, we applied canonical correlation as a spatial filter, which maximizes the covariance between single trials and the average trial^39,40^. By segregating the data into bins of chronological trial numbers, we aimed to reflect experiments under varying measurement time constraints. In a two-factorial repeated measures ANOVA (3×9: measurement modality x trial count), measurement modalities differed significantly in SNR (*F*_(1.89,37.88)_ = 13.13; *p* < .001) and SNRs in general increased with trial numbers (*F*_(1.36,27.24)_ = 47.04; *p* < .001).

The magnitude of SNR increases with trial numbers also differed between measurement modalities (Interaction trial number x measurement modality: *F*_(3.25,64.93)_ = 8.71; *p* < .01; see Fig. 2A). Pairwise post-hoc tests showed that the SNRs of OPM-signals significantly exceeded those of EEG-signals under all trial count conditions. After 20 trials, OPMs reached an average SNR, which was higher than the average SNR of EEG after 80 trials and of SQUID after 40 trials. Overall, there was an increase in SNR for OPMs of up to 205% compared to EEG and of up to 40% compared to SQUID. The SNR of the OPM signal was significantly higher than that of the SQUID signal for 20, 50, 60, 70 and 80 trials. In addition, the SNR of SQUID-signals were characterized by significantly higher SNR compared to EEG-signals (for all apart from 1 and 20 trials). In general, maximum SNRs for OPM-signals were 38% higher than for SQUID-signals and 210% higher than for EEG-signals (see Fig. 2B).

**Figure 2.**
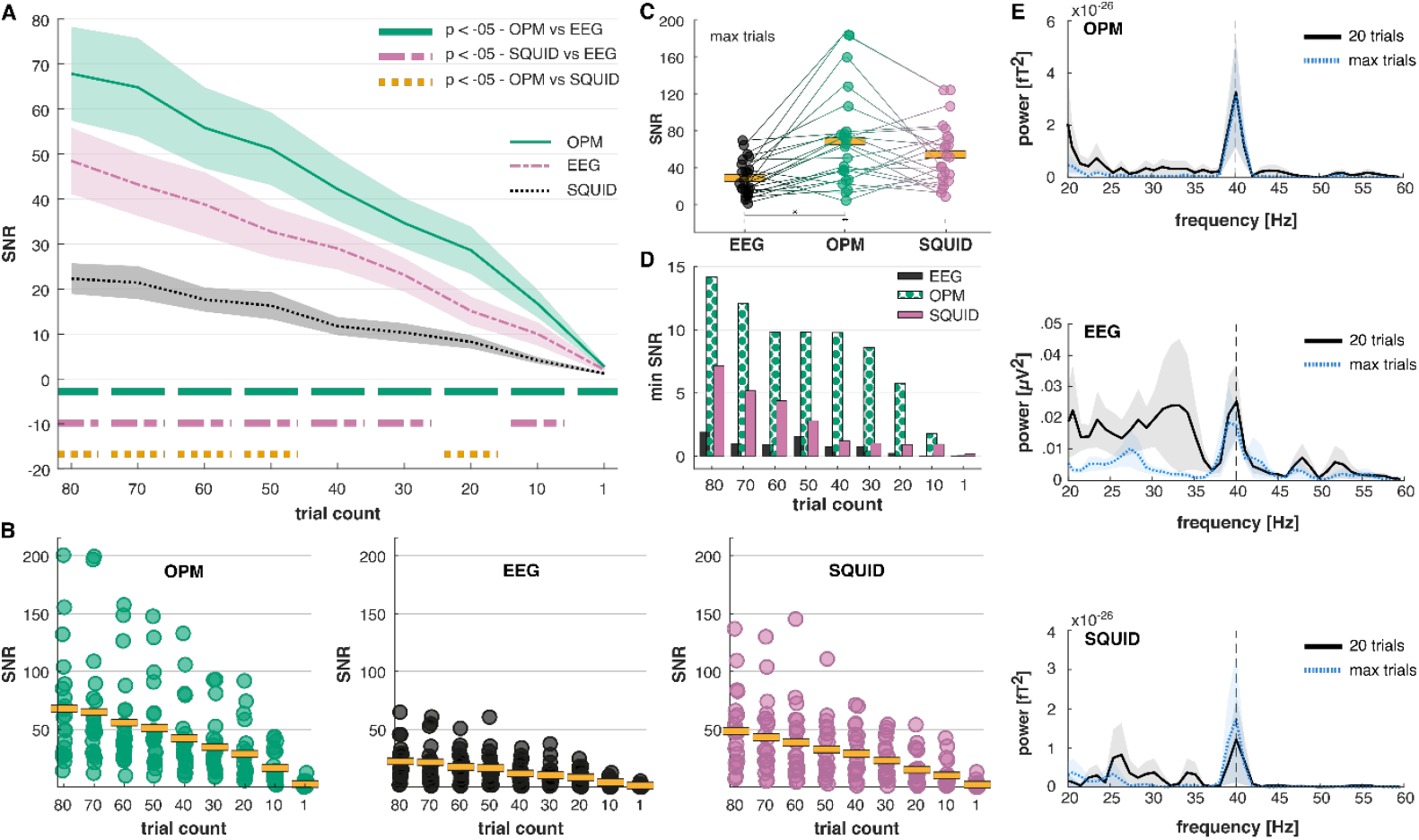
OPMs have significantly higher SNRs compared to EEG and SQUID. **A)** Signal-to-noise ratios (SNRs) of OPMs over trial numbers. The SNRs of OPMs were significantly higher than those of EEG and SQUID. Even when comparing only the first trial of each individual, OPMs have a significantly higher SNR compared to EEG (n = 21). **B)** Illustration of individual SNR values for chronological trial numbers compared between OPM, EEG and SQUID (n = 21). **C)** When incorporating all available trials, OPMs had on average the highest SNRs, significantly higher than EEG (n = 23). **D)** It is notable, that the minimal SNR between measurement condition is consistently highest for OPM, higher by a factor of ∼5-6 for EEG and ∼2 for SQUID (n = 21). **E)** The power spectrum of the trial average compared between 20 trials and the maximum number of trials for each measurement condition. Noise in surrounding frequencies is reduced for OPMs compared to EEG and SQUID, especially with a trial limitation of 20 (n = 21).

In a separate analysis, we allowed trial numbers to differ between measurement modalities, by choosing the maximum trial number after preprocessing. Here, SNRs of both OPM- and SQUID-signals were significantly higher than SNRs of EEG-signals (*F*_(2,44)_ = 11.49; *p* < .001; see Fig. 2C). It is noteworthy that on average (over trial numbers 10 – 80), the minimum SNR was 333% higher for the OPM-signal compared to the SQUID-signal and 2078% higher than the minimal SNR for EEG-signals (Fig. 2D). The difference between measurement modalities was sufficiently salient to be visible by manual inspection of the power-spectra, especially when comparing low trial numbers to maximum trial numbers (see Fig. 2D). Our findings underline the superior measurement precision provided by the OPM-system compared to the EEG- and SQUID-system.

### The phase coherence of OPMs exceeds EEG and SQUID

In addition to measurement precision, we were also interested in the assessment of the coherence across trials compared between EEG, OPM and SQUID. To this end, we calculated a ratio of inter-trial phase (ITPCR) coherence at 40 Hz relative to adjacent frequency-bands during the phase of auditory steady-state stimulation. As ITPC reflects the uniformity of phase-angles in neural responses across trials^41^, it is affected by individual differences. Within individuals, however, it is characterized by high test-retest reliabilities^42–44^. Accordingly, ITPC ratios in our study reflect the coherence with which the measured brain signal is measured rather than actual neuronal differences, especially for the comparison between EEG and OPM, which were recorded simultaneously.

The ITPCR differed significantly between measurement modalities (see Fig. 3A-B; *F*_(2,40)_ = 16.33; *p* < .001) and increased with trial numbers (*F*_(1.75,35.01)_ = 88.67; *p* < .001). This increase, however, was unequal between measurement modalities (*F*_(4.01,80.28)_ = 5.87; *p* < .001). Post-hoc comparisons revealed that the ITPCR of OPM-signals was significantly higher than the ITPCR of EEG-signals across all trial numbers. Overall, the increase in ITPCR for OPMs was up to 61% compared to EEG and 23 % compared to SQUID-systems. Individual ITPCR data points revealed that maximum ITPCRs were comparable between OPM and SQUID (up to 7% higher for OPMs) and up to 52% higher for OPM compared to EEG-signals (see Fig. 3B). Incorporating the maximum trial number for each measurement modality led to significantly higher ITPCRs for OPM- and SQUID-signals compared to EEG-signals (see Fig. 3C; *F*_(2,43.95)_ = 9.77; *p* < .001). The minimum ITPCR for OPM-signals was higher compared to EEG- and SQUID-signals with on average, 76% increase in ITPCR compared to EEG and 51% increase in ITPCR compared to SQUID (see Fig. 3D). These differences are apparent even under manual inspection when comparing the ITPC between 20 trials and the maximum trial number across measurement modalities (see Fig. 3E). Taken together, our results indicate a stronger coherence in measuring neural responses across trials for OPM-systems compared to EEG- and SQUID-systems.

**Figure 3.**
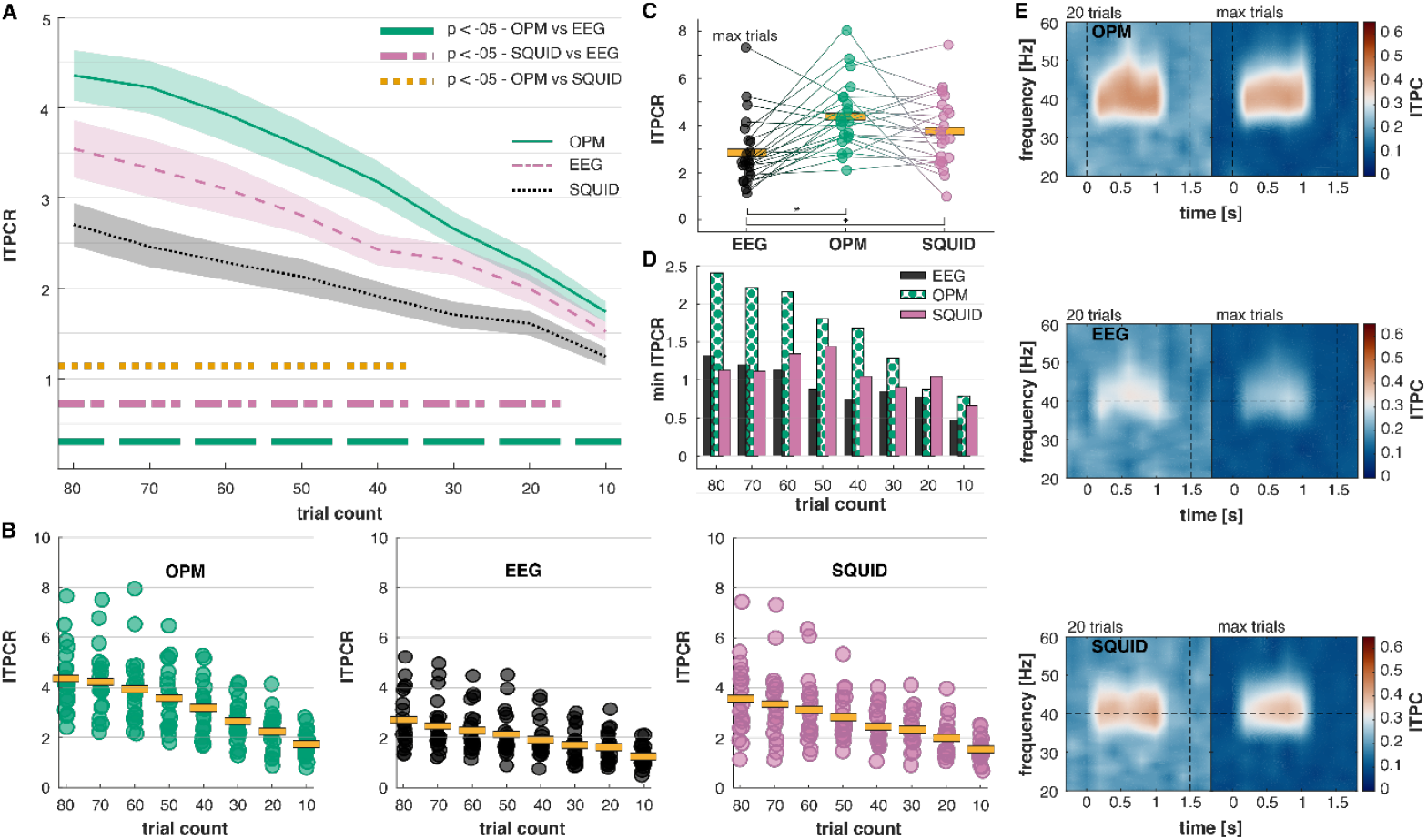
OPMs have significantly higher coherence ratios compared to EEG and SQUID. **A)** Coherence ratios (ITPCRs) of OPMs over trail numbers. The ITPCRs of OPMs were significantly higher than those of EEG and SQUID. This difference between OPMs and EEG was already significant after 10 trials (n = 21). **B)** Illustration of individual CR values for chronological trial numbers compared between OPM, EEG and SQUID (n = 21=. **C)** When incorporating all available trials, OPMs had on average the highest CRs, significantly higher than EEG (n = 23). **D)** It is notable that the minimal ITPCR is highest for OPM, and increased by up to ∼100% compared to EEG (n = 21). **E)** Inter-trial phase coherence compared between 20 trials and the maximum number of trials for each measurement condition. OPMs show the strongest coherence at 40 Hz, even when limiting trial numbers to 20 (n = 21).

### 10-16 OPM sensors versus 64 EEG-Electrodes versus 124 SQUID-gradiometers

As our previous analyses incorporated 10 sensors/electrodes per measurement modality, we aimed to investigate how 10-16 OPMs compared to whole-head EEG and SQUID-MEG measurements. We therefore conducted two separate two-factorial repeated measures ANOVAs (2×9, sensor count x trial count) for EEG and SQUID. We observed a significant increase in SNR with whole-head solution, both for EEG (*F*_(1,20)_ = 56.30; *p* < .001) as well as SQUID (see Fig. 4A; *F*_(1,20)_ = 19.79; *p* < .001). Increases in SNR were different across trial counts (EEG: *F*_(1.82,36.30)_ = 39.79; *p* < .001; SQUID: *F*_(1.99,39.79)_ = 47.22; *p* < .001), with significant increases present in all conditions apart from single trials for EEG and single and 10 trials for SQUID. Additionally, SNRs increased faster with accumulating trial numbers for whole-head-systems compared to limited sensor/electrode setups (Interaction condition x trial count; EEG: *F*_(4.01,80.24)_ = 14.11; *p* < .001; SQUID: *F*_(2.21,44.24)_ = 4.57; *p* = .013).

**Figure 4.**
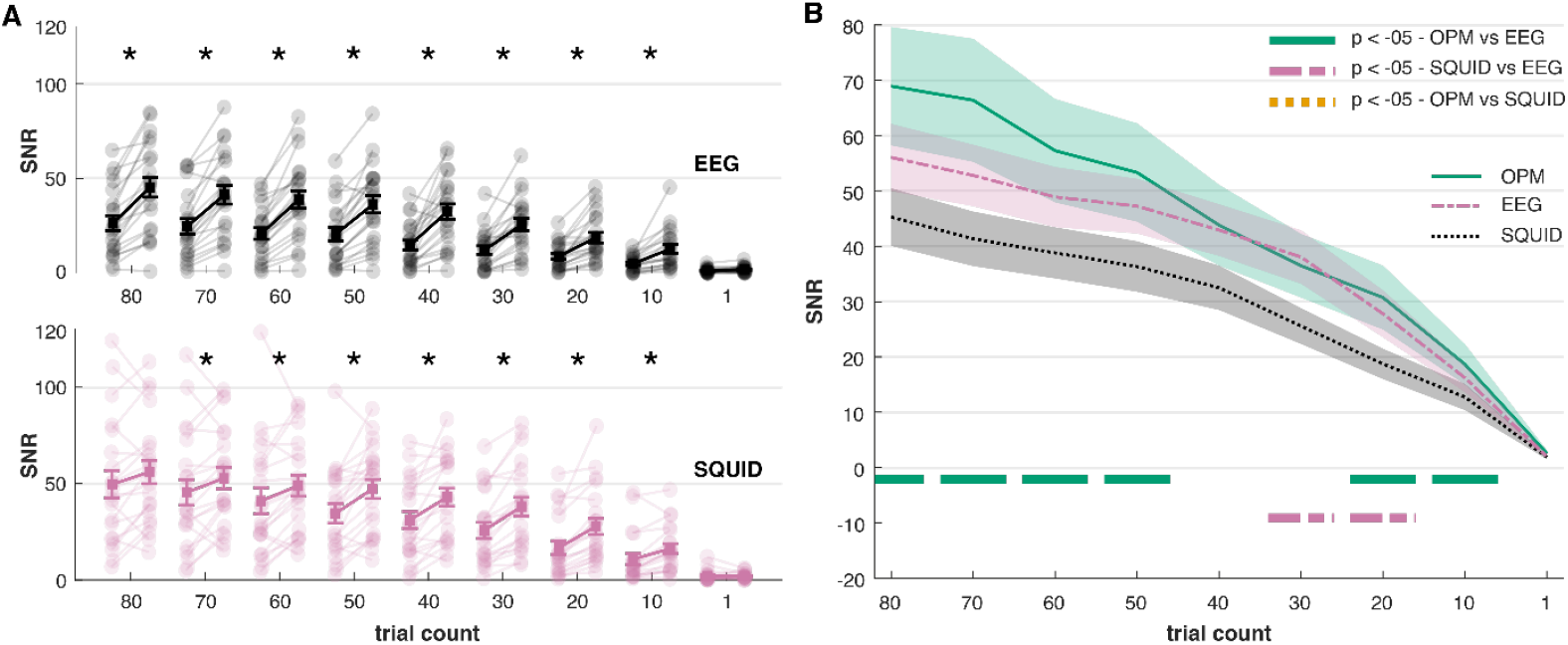
SNR increase from limited sensors to whole-head setup. **A)** For both EEG and SQUID-MEG, there is a significant increase in SNR from 10 sensors/electrodes to a whole-head system (56 EEG electrodes and 125 SQUID gradiometers). Apart from single trials (and 80 trials for SQUID), this difference is significant across trial counts. Post-hoc contrasts with a significance level of p < .05 are marked with * (n = 21). **B)** The SNRs measured with 10-16 OPMs significantly outperform those measured with 56 EEG electrodes for all trial counts but 1,30 and 40 (n = 21). Compared to 125 SQUID gradiometers, the SNRs of 10-16 OPMs are comparable or better, however this difference is not significant.

We then compared SNR values between 10-16 OPM sensors and whole-head EEG and SQUID-systems. Again, measurement modalities differed significantly in their SNR (*F*_(1.79,35.78)_ = 4.01; *p* = .031) and SNRs generally increased with trial count (see Fig. 4B; *F*_(1.63,32.54)_ = 47.94; *p* < .001). The slope of the SNR increase with higher trial counts differed between measurement modalities, as shown in a significant interaction between SNRs for measurement modalities and trial count (*F*_(4.32,86.43)_ = 2.79; *p* < .028). Post-hoc comparisons revealed that SNRs of OPM-signals were significantly higher than those of EEG-signals, for all trial counts apart from single trials and 30-40 trials. Our findings underline the precision with which OPM-sensors measure brain activity, which is elevated in comparison to a whole-head EEG system and comparable to a whole-head SQUID system, even with a limited amount of OPM sensors.

## Discussion

Demands for improvements in non-invasive neuroimaging techniques are rising, as advances in understanding the human brain are hindered by a lack of sensitive measurement instruments and considerable differences in the neuronal structure of model organisms compared to humans^1^.

To investigate whether the latest generation of quantum sensors can push the boundaries of electrophysiological measurements in humans, we assessed auditory steady-state responses with optically pumped magnetometers (OPMs), EEG and conventional MEG (SQUID) within healthy participants (n = 23). So far, there has been no comprehensive statistical validation on the performance of OPM, EEG and SQUID within the same participants. Even though some studies indicated superior performance for OPMs compared to EEG and comparable performance to SQUID, findings were based on small sample sizes and evidence to the contrary has also been reported^2,3,24,31–34^.

Our findings demonstrate that OPMs, a new generation of quantum sensors, surpass EEG and conventional (SQUID) MEG in signal-to-noise ratio (SNR) and coherence across trials of neural responses. SNRs of OPM-signals were up to 200% higher compared to EEG-signals and up to 40% higher compared to SQUID-signals. Additionally, the coherence across trials was equally higher for OPMs, with up to 40% increase compared to EEG and up to 20% increase compared to SQUID. Our findings emphasise the potential of OPMs to advance non-invasive human neuroimaging, as previous studies demonstrate the paramount importance of increased SNR in enhancing the reliability and sensitivity of numerous methods and analyses, including the accurate estimation of functional connectivity, source reconstructions, and diagnostic markers^45–47^.

This is important, as accumulating data suggests a need for higher differentiation between neuronal circuits, which have been shown to be functionally and locally distinct, despite producing similar electric potentials^48,49^. In addition, we showed that OPMs provide significantly higher measurement coherence than EEG and SQUID, which relates to increased uniformity of phase angles across trials. Measurement coherence can be affected by noise, which may mask the phase angles of interest, and by a mixing of sources. This is most prevalent in EEG, where different neuronal circuits produce measurable activity, conducted through brain tissue and mixed on the scalp^12,13^. Accordingly, the strength with which a circuit is represented on the scalp signal in one trial, may not be the same in another, ultimately resulting in slight latency differences and reduced coherence measurements. Enhanced measurement coherence in OPMs compared to EEG, therefore points to increased consistency in the acquisition of neuronal circuit activity. In fact, there are simulations which propose the possibility of OPMs to differentiate between neuronal signals in higher compared to lower layers of the cortex^5^. The increased coherence across trials for OPMs compared to SQUID-sensors may likewise be caused by increased consistency in the measurement of neural circuit activity. As trial coherence is not independent of amplitude, however, an alternative explanation could stem from increased signal amplitudes for OPM^3^ compared to SQUID-measurements, resulting from the reduced distance of the sensors to the human scalp.

A prior study investigated evoked potentials, but unlike in our results, the authors found equivalent or lower SNRs for OPM-compared to SQUID-signals^29^. These discrepancies may be explained by sensor-to-signal distance, which affects SNR, especially when sensor density is unequal. To prevent this bias, we matched sensor numbers and positions between systems and applied a spatial filter^50^, to maximize the representation of the signal of interest across sensors. Consistent with the previous study, we found no SNR differences across modalities when using only the highest-SNR sensor/electrode, which generally led to strongly reduced SNRs, indicating that single sensors do not exhaust the full potential of each measurement method (see SUPPL Fig. 1).

Importantly, we showed that after 20 trials, OPM-signals reached average signal-to-noise ratios (SNRs), which required 80 trials with EEG and 40 trials with SQUID. Even on the single-trial level, the difference in SNR was significant. These findings introduce two possibilities: to increase the complexity of task designs in basic neuroscience and to reduce required measurement time in translational or clinical neuroscience, which is especially relevant for paediatric or clinical populations (with difficulties to endure lengthy measurement sessions, such as infants or chronic pain patients)^7,51^. This is further underlined by the fact, that minimal SNRs for OPMs were on average 2078% higher than for EEG and 333% higher than for SQUIDs, showing that in recordings where noise was relatively high, OPMs were better at capturing the signal of interest. Furthermore, increased SNR for single trials improves the application of brain-computer interfaces, which have gained importance in clinical application, such as for brain-state dependent neuromodulation^6^. In fact, recent studies indicated improved performance for an OPM-based brain-computer interface compared to SQUID^20,52^. Additionally, we showed that 10-16 OPM sensors were sufficient to significantly surpass SNRs acquired with a whole-head EEG and achieve on average higher SNRs acquired with a whole-head SQUID system. This pattern has been demonstrated previously, however without sufficient data to perform statistical validation^2,3,15^.

It should be noted that there are certain caveats in comparing different measurement modalities. Neuronal responses to auditory stimulation in general may be more easily recorded with OPM and SQUID systems compared to EEG, based on the source orientation in Heschl’s Gyrus^40,44,53^. Furthermore, the SQUID system in this study comprised gradiometers, which improve SNRs by reduction of non-neuronal noise^54^. The OPM-sensors applied in this study, on the other hand, comprised two measurement directions each, which increases channel density, benefitting the application of spatial filters, such as canonical correlation^50,55^. The EEG-OPM recording was performed upright, while the SQUID recording was performed in supine position. There are differences in neuronal signal strength between these positions, however, it is unclear how auditory steady-state responses are affected^56–58^. Lastly, it is noteworthy that the SQUID-recording took place in a room with reduced shielding compared to the EEG-OPM recording. However, due to the fact that SQUID-MEG systems are static and therefore not affected by gradients in the background fields, the applied shielding is sufficient to suppress environmental noise in our setting^59^.

Taken together, our findings demonstrate a significant increase in SNR and coherence across trials for OPMs compared to the current gold-standards of electrophysiology. These improvements directly impact the measurement quality of neural circuit activity, while also reducing the time required for recordings. Consequently, OPMs have the potential to significantly advance neuroscience research, brain-computer interfaces and precision diagnostics and therapy.

## Methods

### Participants

We recruited N = 23 healthy volunteers between 18 and 55 years (mean = 36 years ± 10; 11 = female). Participants were recruited via university mailing lists and online advertising. At the end of the experiment, participants received monetary compensation (10€ /h). The study protocol was approved by the Ethics Committee of the Charité University Hospital, Berlin and in accordance with the Declaration of Helsinki. All participants provided written informed consent.

### Data Acquisition

EEG and OPM-MEG measurements were performed using a modified 56-electrode cap (equidistant layout) by ANT Neuro with tailor-made OPM holders (see Fig. 1 A). Participants were seated inside an 8+1-layer magnetically shielded room at Physikalisch-Technische Bundesanstalt (PTB), Berlin^60^. Additional AgCL ring electrodes were placed next to the eye (electrooculogram) and at the left and right earlobe (reference and ground). Impedances were kept below 10 kΩ.

10-16 dual-channel OPM sensors (QuSpin QZFM 2^nd^ generation) were positioned over left and right temporal regions (see Fig. 2). Online OPM-MEG recordings were sampled at 4000 Hz and EEG recordings were sampled at 500 Hz. Visual stimuli were presented via a shielded projector^61^.

SQUID-MEG data was acquired in a separate session using a Yokogawa system with 124 gradiometers. This session of the experiment took part in a 2+1 layer shielded room. Participants were lying in supine position, while visual stimuli were projected onto a screen. Online SQUID-MEG recordings were sampled at 4000 Hz. To avoid confounds due to repetition and learning effects, half of the participants performed the EEG-OPM session first, while the other half started with the SQUID-MEG session.

For all measurements, auditory stimulation was applied utilizing an etymotic® sound system and transmitted into the shielded room via tubes, which were connected to in-ear plugs (Doc’s Promolds).

### Paradigm

Prior to recordings, individual hearing thresholds were assessed utilizing a staircase procedure. Participants focused on a central fixation cross while sounds were played in the background which consisted of 1000 Hz carrier tones, which were either amplitude modulated at 40 Hz (100 trials), at 20 Hz (4 trials) or at 60 Hz (4 trials), presented in random order (see Fig. 1C). Trials modulated at 20 Hz and 60 Hz were accompanied by arrows either pointing to the left or to the right. Participants were asked to press a button when arrows pointing to the right appear, and to ignore arrows pointing to the left. Stimulation was performed via PsychoPy^62^ for the EEG-OPM recordings and with Presentation® software (Version 21.0, Neurobehavioral Systems, Inc., Berkeley, CA, www.neurobs.com) for the SQUID-MEG recordings.

### Data Preprocessing

The preprocessing steps were identical across measurement modalities. Data was segmented into 4s epochs starting 1s prior to ASSR stimulation and down sampled to 500 Hz. EEG data was re-referenced to an average reference. To remove line-noise, data was notch-filtered between 49 and 51 Hz. For visual artifact inspection, the dataset was filtered between a) 110 and 140 Hz and to manually identify trials with strong muscle artifacts and between b) 0.5 and 80 Hz to identify other movement artifacts, as well as noisy or flat channels. Additionally, a semi-automatic algorithm was used to visually identify trials with strong deviations in amplitude variance (see Table 1 for number of removed trials/channels/components per modality).

**Table 1.** average removed trials, channels and ICA components per modality.

|  | EEG | OPM-MEG | SQUID-MEG |
| --- | --- | --- | --- |
| trials | 18 ($\pm 17$) | 20 ($\pm 21$) | 9 ($\pm 11$) |
| ICA components | 3 ($\pm 1$) | 3 ($\pm 2$) | 3 ($\pm 1$) |
| channels | 0 ($\pm 1$) | 1 ($\pm 2$) | 3 ($\pm 1$) |

*note: numbers are presented as mean ( ± SD)*

As the amount of removed trials directly impacts signal-to-noise ratios, it could be argued that removed trials should be identical between EEG and OPM-MEG. However, as the measurement modality may directly impact the number of trials that need to be removed and could therefore be considered an important factor in the evaluation of the method, we decided to preprocess EEG and OPM-MEG data separately (see Table 1 for number of removed trials/channels/components per modality).

The data was filtered between 5 and 40 Hz to identify eye-blink-, eye-movement- and heartbeat-components using an independent component analysis (ICA). Finally, components were removed and the cleaned dataset was filtered between 5 and 60 Hz.

To retrieve channel weights which optimally represent the measured signal, a spatial filter was created using canonical correlation^50,55^. To this end, channel weights which maximized the covariance between the bandpass filtered single trials with the bandpass filtered average signal during the time of stimulation (0.25s – 1s) were calculated for each individual. The channel weights were then applied to the dataset filtered between 5 and 60 Hz. As canonical correlation is insensitive to polarity, an algorithm was devised which identified local minima and maxima of the average over all participants and channels (without canonical correlation). These datapoints were then applied to individual datasets. If the average over all local minima was higher than the average over all local maxima, the data was multiplied by -1. Afterwards, the average signal was inspected manually per participant to confirm the polarity based on event-related fields/potentials. EEG electrodes were chosen based on the maximum measured activity for 40 Hz and SQUID-MEG gradiometers were chosen based on their topographic proximity to the positions of OPM sensors (see Fig. 2D for layouts of all modalities and the chosen electrodes/sensors). All preprocessing steps and analyses were performed in MATLAB R2020b and the fieldtrip toolbox version 20221223^63^.

### Data Analysis

To assess SNR between measurement modalities, spectral power of the trial average and inter-trial phase coherence (ITPC) were computed. Spectral power was calculated between 0.25 and 1s using a sliding hanning-taper with a dynamic duration of 7 cycles (frequency range: 1 to 60 Hz, step size: 0.5 Hz). ITPC was calculated using a sliding gaussian wavelet with a dynamic duration of 7 cycles. The frequency ranged from 1 to 60 Hz in a step size of 0.1 Hz and the temporal resolution was set to 0.25 between -1s and 2s.

SNRs were calculated by comparing 40 Hz spectral power and ITPC with adjacent frequencies. As such, power SNRs constituted a comparison between 40 Hz and the mean across a noise frequency-range between 20 and 38 Hz as well as 42 Hz – 48 Hz, which was normalized applying the following formula: 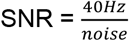. For ITPC-data, the noise frequency range was limited to 20-30 Hz to avoid frequency-smearing of signal into the noise range.

To compare SNR estimates for different trial numbers, we successively reduced the number of trials inserted into the power/ITPC estimation used for SNR-calculation. Trials were always chosen in chronological order to best represent real experiments with only for instance 10 trials. For this analysis, we removed 2 participants as both had less than 70 trials left after preprocessing in at least one measurement condition. Percent increase between SNRs of measurement modalities was calculated as subtraction of the increased value from the original value, divided by the original value and multiplied by 100.

Statistical differences between modalities were assessed using repeated-measures ANOVA utilizing Tukey’s Honest Significant Difference as post-hoc assessment. To assess post-hoc contrasts over different trial numbers, separate two-factorial repeated measures ANOVAS between pairwise measurement modalities were calculated. Sphericity of the factor variables was investigated with Mauchly’s test. Cases which violated the assumption of sphericity were corrected using the Greenhouse Geisser method and adjusted p-values and degrees of freedom are reported.

## Acknowledgements

We thank the DFG for funding this project and ANT-Neuro for cooperating with us in the modification of their EEG-cap. Additionally, we would like to thank our colleague Dr. Tineke Grent-t’-Jong, who provided a lot of theoretical and practical support in the project. Lastly, we would like to express our gratitude to Anna Benedict, Carlotta Preller, Katharina Stammkötter and Fabian Symanowski, who tirelessly helped to acquire all the data for this project.

## Author information

MB has conceived the idea, collected the data, programmed the task, performed the analysis and devised the manuscript. PA has collected the data, programmed the task, created the technical setup and revised the manuscript. PK has provided technical and theoretical support and supervision. TS has conceived the idea and provided technical and theoretical supervision. PJU has conceived the idea and provided supervision.

## Supplementary Information

**SUPPL Figure 1.**
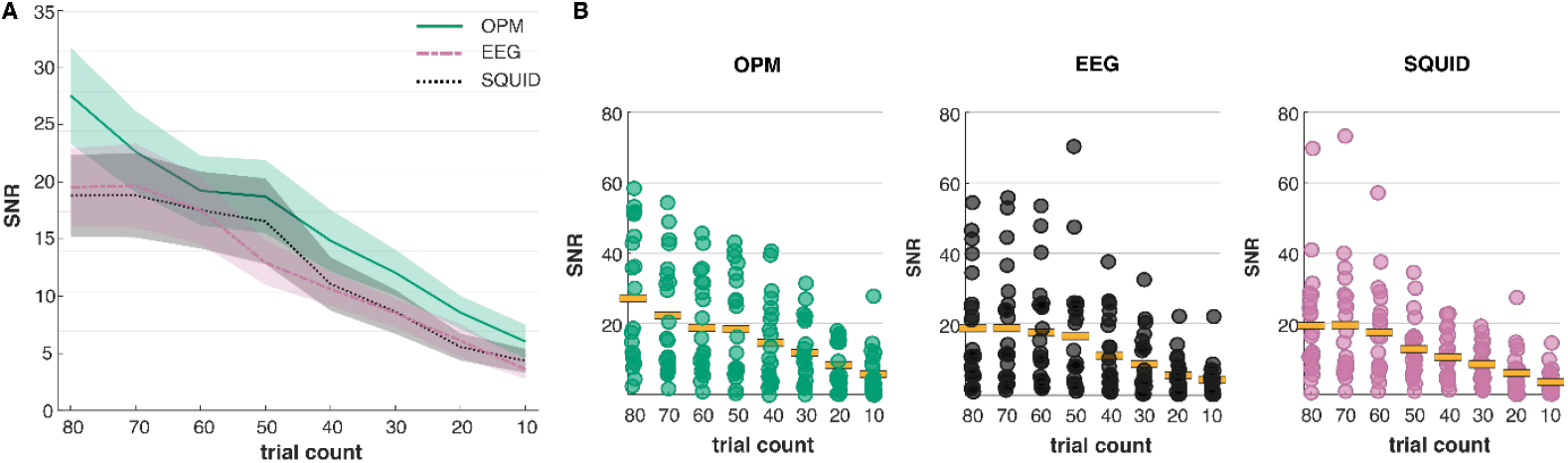
SNR comparison between OPM, EEG and SQUID-systems for single highest SNR sensor/electrode. **A)** On average, OPM-signals show the highest SNR values across trial counts, however, differences between measurement modalities are not significant (n = 21). **B)** A closer look at individual SNR-values illustrates that SNR-values across modalities are comparable when selecting a single sensor with the highest SNR for each individual. SNR-values on average are decreased by a factor of ∼2-3 compared to the application of spatial filters (see Fig. 2).

